# Capturing in-field root system dynamics with the RootTracker

**DOI:** 10.1101/2020.11.14.382002

**Authors:** Jeffrey J. Aguilar, Matt Moore, Logan Johnson, Rachel F. Greenhut, Eric Rogers, Drew Walker, Fletcher O’Neil, Jake L. Edwards, Jake Thystrup, Sam Farrow, Jesse B. Windle, Philip N. Benfey

## Abstract

Optimizing root system architecture offers a promising approach to developing stress tolerant cultivars in the face of climate change, as root systems are critical for water and nutrient uptake as well as mechanical stability. However, breeding for optimal root system architecture has been hindered by the difficulty in measuring root growth in the field. Here, we describe a technology, the RootTracker (RT), which employs capacitance touch sensors to monitor in-field root growth over time. Configured in a cylindrical shutter-like fashion around a planted seed, 264 electrodes are individually charged multiple times over the course of an experiment. Signature changes in the measured capacitance and resistance readings indicate when a root has touched or grown close to an electrode. Using the RootTracker, we have measured root system dynamics of commercial maize hybrids growing in both typical Midwest field conditions and under different irrigation regimes. We observed rapid responses of root growth to water deficits and found evidence for a “priming response” in which an early water deficit causes more and deeper roots to grow at later time periods. There was genotypic variation among hybrid maize lines in their root growth in response to drought, indicating a potential to breed for root systems adapted for different environments.

## Introduction

Yield stability in agriculture is a major challenge in the face of climate change, necessitating crop varieties that are resilient to stressful environmental conditions such as drought. Optimizing root system architecture offers a promising approach to developing stress tolerant cultivars, as root systems are critical for water and nutrient uptake as well as mechanical stability. However, breeding for optimal root system architecture has been hindered by the difficulty in measuring root growth in the field. Current methods are laborious and not easily scaled. To enable the much broader use of root system architecture information in agronomic research, we have developed a device that overcomes these challenges.

Optimization of root system architecture could provide benefits beyond stress tolerance in the face of climate change to actually mitigating climate change by reducing greenhouse gas levels. Roots contribute to carbon sequestration in soil through exudates and cell wall material (Dakora and Phillips 2002, Paul and Clark 1996). An analysis of soil organic carbon in croplands indicates that modulating root growth can significantly impact greenhouse gas emissions mitigation. Larger and deeper root systems can result in greater deposits of organic carbon compounds with longer mean residence times (Kell 2012). Further, there are strong arguments that deeper root systems provide enhanced water and nitrogen uptake in many circumstances (Lynch 2013). Thus, it is possible to optimize root systems for both abiotic stress tolerance and carbon sequestration, simultaneously.

To breed for optimal root systems, one needs to measure root growth in the field. Common methods used for root phenotyping in the field are shovelomics (Trachsel et al. 2011) and coring (Wasson et al. 2014). While advances in image analysis (Das et al. 2015, Colombi et al. 2015) have allowed for higher throughput of shovelomics phenotyping, both shovelomics and coring are destructive and allow for only a single snapshot of root growth. Another option, minirhizotrons, are tube-encased cameras that are inserted and left in the soil, which typically only image a small subset of the root system (Rytter and Rytter 2012). Among noninvasive methods (reviewed in Wasson et.al 2020), ground penetrating radar (Delgado et al. 2017) has been used to measure bulk root properties. However, to date, it has been limited to use with roots of relatively large diameter. Another approach involves charging a circuit with one electrode connected to the plant’s stem and the other to the soil. This technique has correlated capacitance with tree root length (Ellis et al. 2013) and root mass (Dalton 1995, van Beem et al. 1998) of herbaceous crops, but does not provide a measure of growth rates.

Stress tolerance can be enhanced by optimization of root system architecture, as well as root system dynamics - how root growth changes over time and responds to changes in the environment (Arsova et al. 2020). Measuring root systems in 4D (space and time) has been achieved primarily in lab settings with techniques such as X-ray computed tomography, magnetic resonance imaging, and positron emission tomography (Atkinson et al. 2019).

Although current methods for phenotyping roots have limitations, breeding for root traits has been shown to enhance plant performance (Tracy et al. 2019). In rice, steeper (Uga et al. 2011) and deeper (Hurd 1974, Wasson et al. 2014) root systems were selected for water capture in deeper soil strata, and thicker primary root systems were selected for greater biomass in drought-prone areas. Identifying and connecting such phenotypic responses in field conditions to crop performance metrics such as yield stability (Wang, et al. 2014) will aid in developing crop varieties robust to climate change.

Here, we describe a technology, the RootTracker (RT), which employs capacitance touch sensors to monitor in-field root growth over time. We have used the RT platform to measure root system dynamics of commercial maize hybrids growing in both typical Midwest field conditions and under different irrigation regimes in water-controlled environments. In these experiments, we discovered remarkably rapid responses of root growth to water deficits. We found evidence for a “priming response” in which an early water deficit causes more and deeper roots to grow at later time periods. There was genotypic variation among hybrid maize lines in their root growth in response to drought, indicating a potential to breed for root systems adapted for different environments.

## Results

### The RootTracker uses capacitance touch sensing to detect roots

The RT uses a vertical array of 22 equally spaced capacitance sensing electrodes embedded in each of 12 circuit boards (paddles) which are arranged in a cylindrical shutter-like fashion (Fig. 1(a)). Paddles have V-shaped ends to facilitate entry into the soil. The electrodes are connected to electronics on a ring-shaped circuit board which is covered in urethane for mechanical strength and water-proofing. RTs can be installed in field soil (Fig. 1(b)) using hammers or hydraulic presses. Once installed in soil, the electrodes range in depth from 1.9 to 16.1 cm. Seeds can either be sown in the center of the device after installation, or the RT can be centered and inserted over a growing seedling.

**Figure 1.**
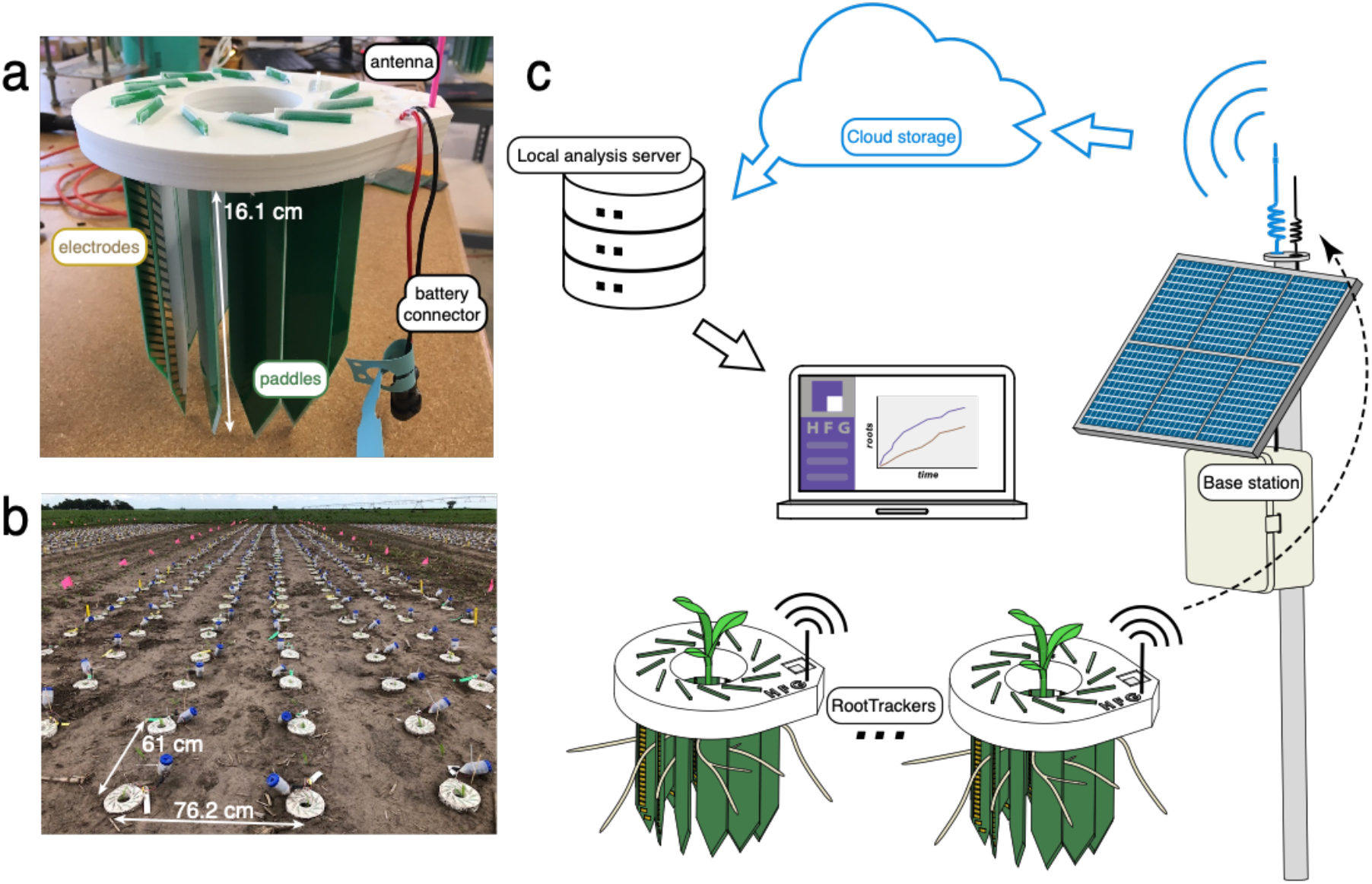
The RootTracker (a) consists of 22 electrodes on each of 12 paddles for detecting roots. Hundreds of RTs in a field (b) communicate raw sensor data via radio transmission to a central base station, which sends the data to cloud-based servers, where the data can be analyzed on local servers for extracting root detections (c).

After the RT is powered (via a 4-AA battery pack), the RT takes multiple raw voltage measurements at each electrode and communicates this data wirelessly to a central base station, which then uploads the data to a cloud-based server (Fig. 1(c)). Once downloaded locally, raw voltage data is processed and converted into electrical capacitance and resistance signals (see Methods for more details). We identified signature fluctuations in the resistance/capacitance space that indicate root growth activity near the sensors once signal changes have been normalized across all electrodes. In this way, fluctuations are evaluated to detect roots. Because all RTs are oriented in the same way and the planting depth is consistent across the field, RT detections capture the initial depth and direction of a plant root’s growth. Since these detections are timestamped, root growth rates can be computed.

### Roots reduce growth in response to water deficit

To examine the response in root growth of different maize varieties to water deficits, we installed 881 RTs in alluvial soil at Massai Agricultural Services in Rancagua, Chile under drip irrigation. We selected 10 genotypes (4 hybrids and 6 inbreds) and subjected them to two watering treatments, well-watered and drought. Fields under the well-watered treatment followed a schedule of periodic irrigation for 53 days (Fig. 2(b)). Fields under the drought treatment had the same irrigation schedule until day 36, when the water was shut off for the remainder of the experiment (a 16-day period).

**Figure 2.**
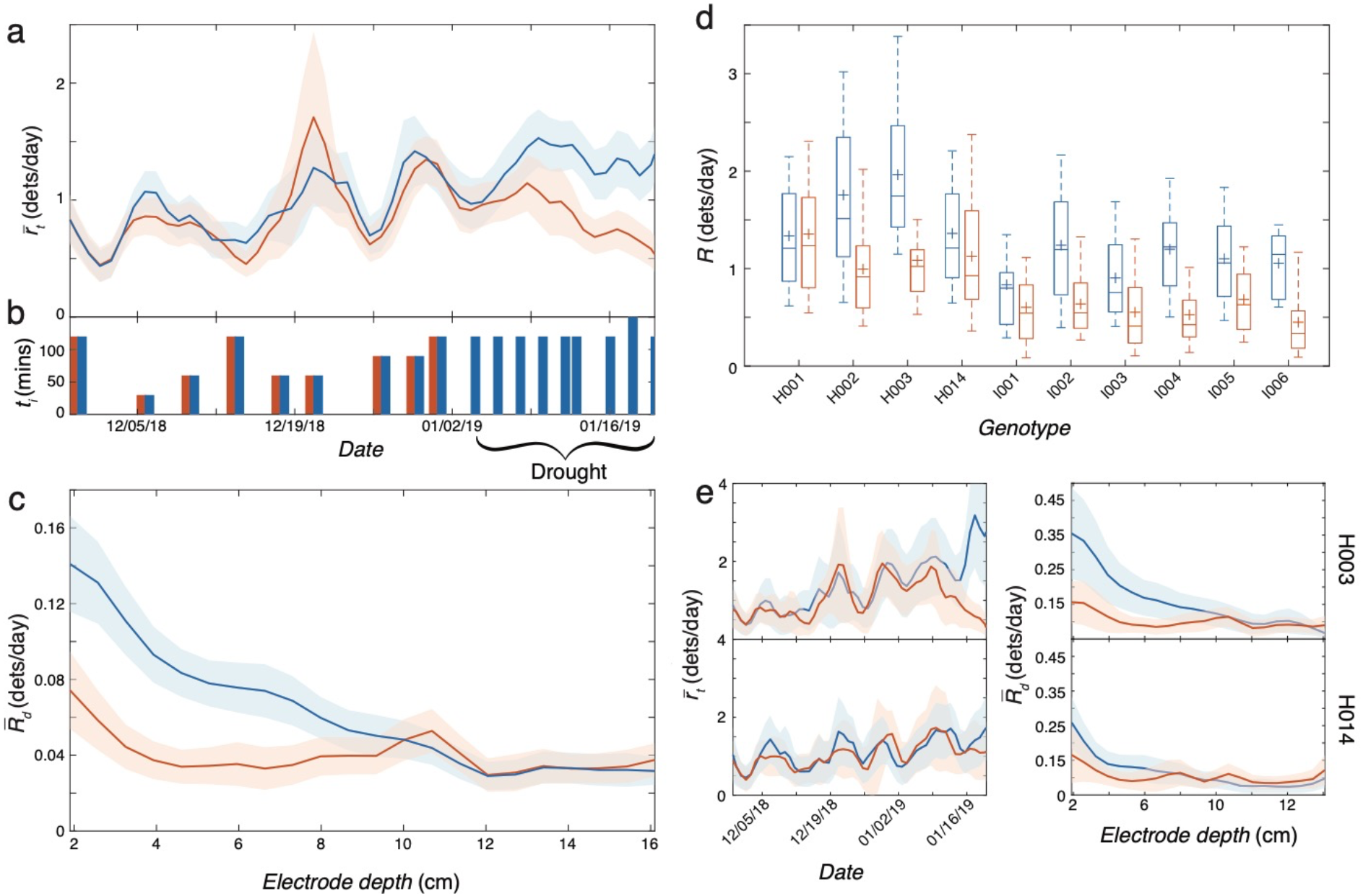
Trial 1: Response to imposed drought. Comparison of root detections between well-watered plants (blue) and drought treatment plants (orange) of (a) smoothed mean daily root detection rate over time, 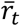, and (c) smoothed mean of the per-electrode-depth daily root detection rate, time-averaged across the drought period (1/4/19 - 1/20/19), 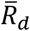. (b) Duration, *t_i_*, of each irrigation event during the trial for each treatment. All shaded regions indicate ± 2 standard errors from the mean. (d) Box and whisker distributions of time-averaged daily root detection rate, *R,* during the drought period, separated by genotype and treatment (blue, well-watered; orange, drought). Top and bottom of box indicate 25 and 75 percentile of RTs, horizontal line in box is median, cross is mean, and whiskers are 9 and 91 percentiles. (e) Well watered and drought treatment comparison of 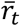 (left) and 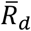 during the drought period (right) for two hybrids (H003, top, and H014, bottom) with variable relative responses to drought.

Grouping all plants by watering treatment, RT detection analysis indicated that plants responded to the imposed water deficit by rapidly reducing root growth. The average daily root detection rate 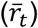 under both treatments was approximately the same until the water deficit was imposed (Fig. 2(a)). The mean time-averaged daily root detection rate, 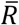, for drought treatment plants during the drought decreased to 0.81 root detections per day - 36% less than the well-watered plants during the same time period (T-test, *p*=<2.2e-16). When analyzed by electrode depth, drought treatment plants exhibited lower average daily growth rates than well-watered plants during the drought in the shallow soil strata, while at deeper soil levels there was little difference in the number of roots detected (Fig. 2(c)). These results indicate fewer roots grew near the surface, likely owing to drying of the soil surface after the irrigation was shut off. During this time, deeper root detection rates remained the same between the two treatments.

To examine genotypic differences in response to the water deficit, we analyzed root growth of each genotype separately (Fig. 2(d)). While we found no significant interaction effect among inbred genotypes, comparing daily detection rates time-averaged during the drought period revealed a significant interaction effect of hybrid and watering treatment (Interaction 2-way ANOVA, *p*=0.003), indicating variation in drought response by hybrid. Two genotypes (H003 and H014, Fig. 2(e)) highlight this divergent behavior. Interestingly, during the drought, H014 plants in both watering treatments as well as H003 plants in the drought treatment all exhibited similar daily root detection rates. Thus, rather than having a decreased negative response to drought, H014 seems to have exhibited a decreased positive response to receiving water.

### Root system dynamics may contribute to priming

Mild drought stress early in the growth of wheat can cause a “priming effect”, which appears to allow plants to better tolerate later episodes of reduced water availability (Wang, et al. 2014). If a similar priming effect occurs in maize, we hypothesized that root system dynamics could play a role in this response. To test this hypothesis, we installed 721 RTs at Massai in Rancagua, Chile divided among 12 hybrid genotypes and two irrigation treatments: well-watered, and water-limited. In the latter irrigation treatment, we imposed two separate water deficits: the first starting 9 days after planting and lasting 6 days, the second starting 35 days after planting and lasting 11 days (Fig. 3(d)).

**Figure 3.**
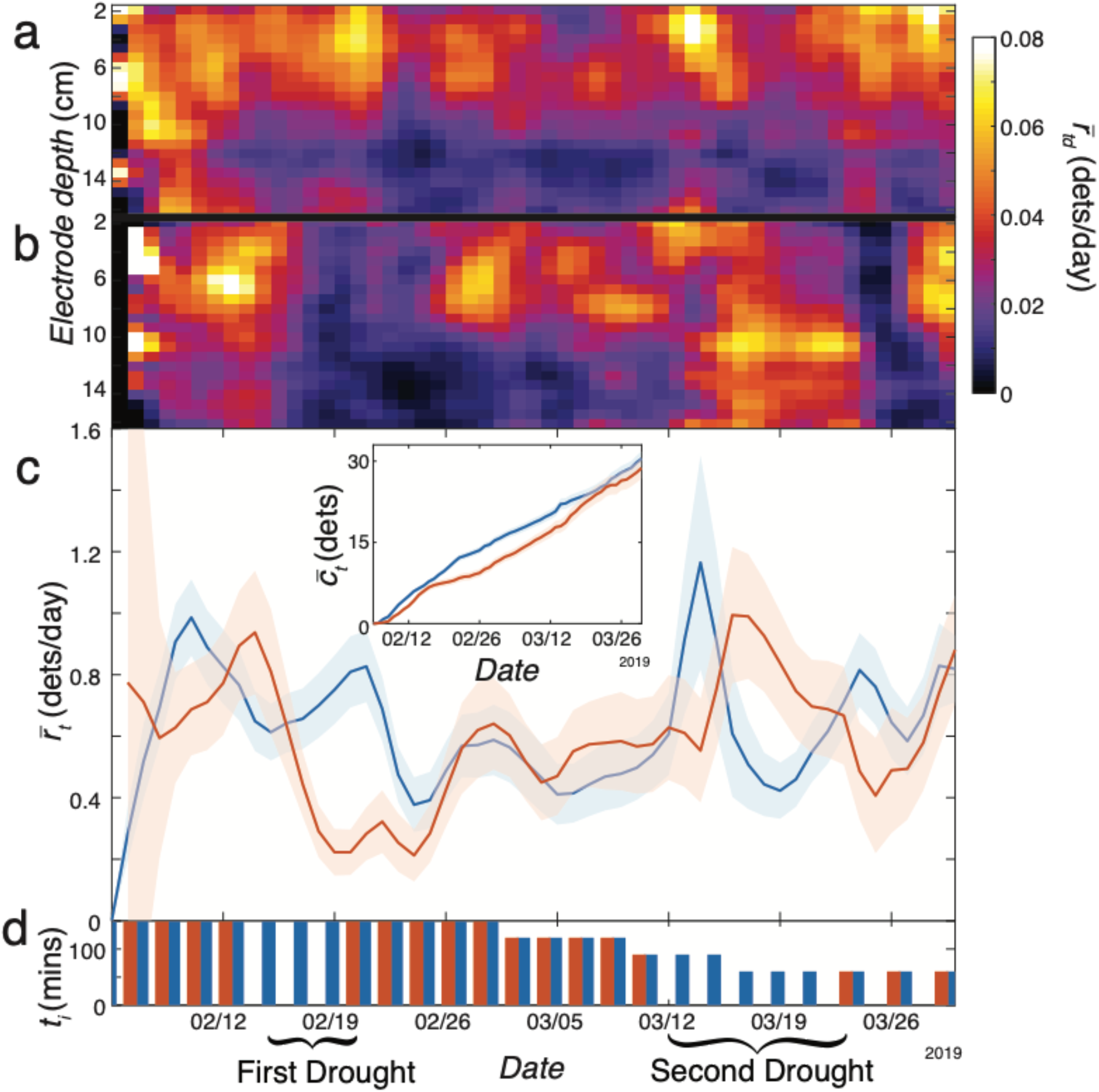
Trial 2: Priming response to early drought. Comparison between well-watered plants (a) and water-limited plants (b), of smoothed mean daily root detection rate over time and electrode depth, 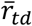. See Supplemental Figure 7 for complementary heat maps of standard error. Comparison between well-watered plants (blue) and water-limited plants (orange) of (c) smoothed mean daily root detection rate over time, 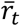, and (c inset) mean cumulative root detections over time, 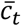. All shaded regions indicate ± 2 standard errors from the mean. (d) Duration, *t_i_*, of each irrigation event during the trial for each treatment.

During the first drought period, we observed a rapid decrease in root growth rate, similar to Trial 1 (Fig. 3(c)). Remarkably, during most of the second drought, root growth was greater in water-limited plants than plants in the well-watered treatment. The average daily rate of detection between 3/15/19 and 3/22/19 for water-limited plants was 0.88 roots per day – 148% higher than well-watered plants (T-test, *p* = 7.84e-13). Comparison of cumulative root growth over time suggests that the subsequent increased root detections allowed these plants to approach the average cumulative detections of well-watered plants (Fig. 3(c) inset). Additionally, during the same time period that we observed overall increased water-limited root growth, water-limited root detections were concentrated at the deeper electrodes, whereas the well-watered root detections were concentrated at more shallow electrodes (Fig. 3(a-b)). From 3/15/19 to 3/22/19, we observed a significant increase in average daily growth rate at the deeper half of electrodes in water-limited plants as compared to well-watered plants (T-test, *p*< 2.2e-16). The mean growth rate in the deep half in water-limited plants was 0.50 roots per day – 319% greater – than that in well-watered plants. These results indicate that an early water deficit can promote more root growth at deeper soil strata later in the growing season, even during a second imposed drought.

### The priming response is independent of a second water deficit

To determine if a second water deficit is required to induce the increased root growth observed in Trial 2, we conducted a follow-up experiment at the Kearney Agricultural Research and Extension (KARE) Center located in Parlier, California during the summer of 2019. Soil and climate conditions resembled those found at the test site in Chile. We used 1457 RTs divided among 10 hybrid genotypes (six of which were included in Trial 2) and three irrigation treatments. The well-watered treatment followed the irrigation schedule illustrated in Fig. 4(b). The furrows between the planted rows were drip irrigated to produce consistent water diffusion in the soil. The single drought treatment consisted only of an early water deficit (14-day water shutoff starting 16 days after planting). The double drought treatment consisted of the same early drought, as well as a second drought (16-day water shut-off starting 44 days after planting), similar to the water-limited treatment in Trial 2.

**Figure 4.**
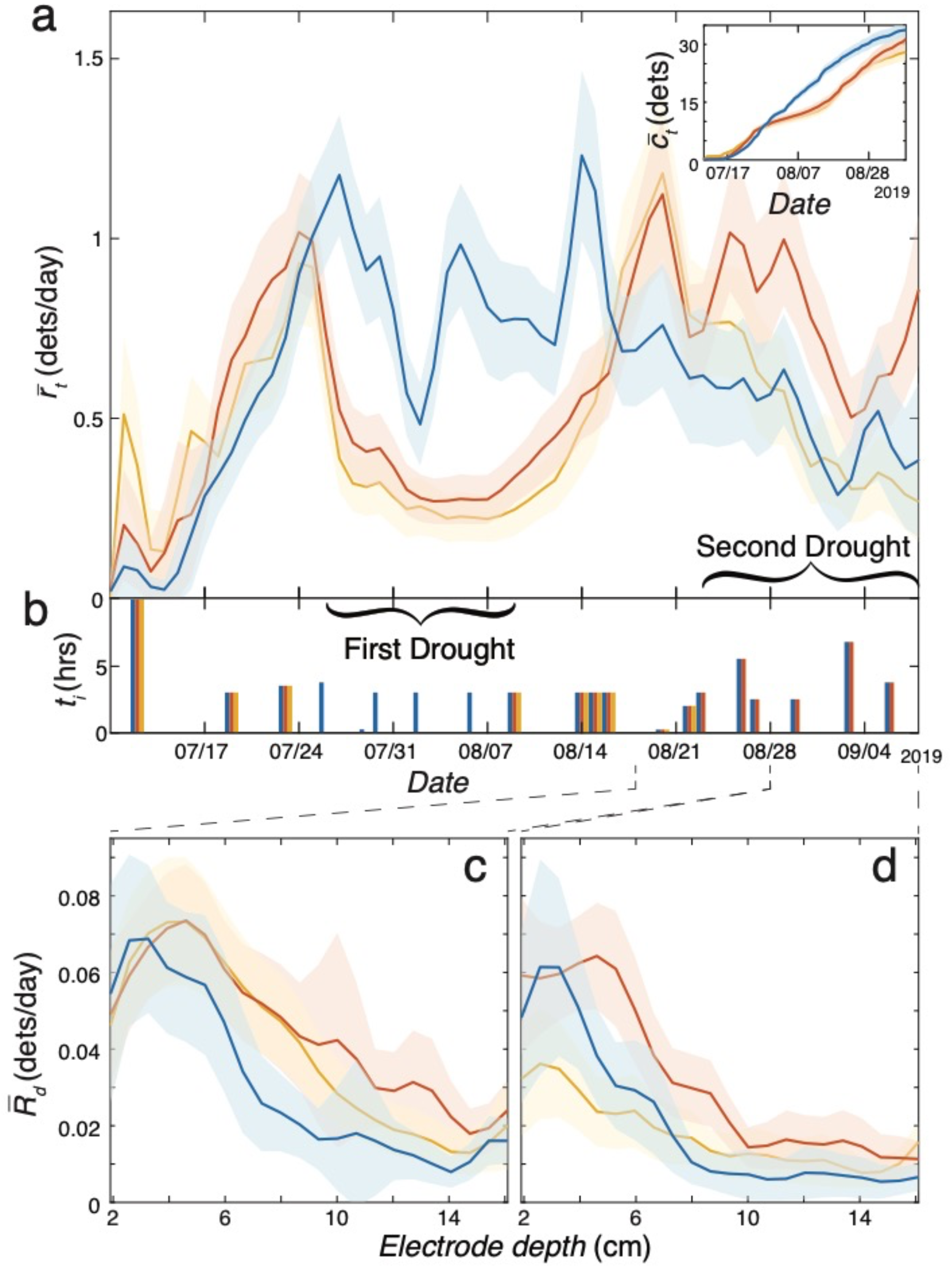
Trial 3: Priming response to early drought. Comparison between well-watered plants, (blue), single drought plants, orange, and double drought, yellow, of (a) smoothed mean daily root detection rate over time, 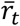, (inset), mean cumulative root detections over time, 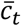, and smoothed mean of the time-averaged, per-electrode-depth daily root detection rate, 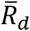, from (c) 8/18/19 to 8/28/19 and (d) 8/28/19 to 9/8/19. All shaded regions indicate ± 2 standard errors from the mean. (b) Duration, *t_i_*, of each irrigation event during the trial for each treatment. See Supplemental Figure 6 for per-pedigree and treatment distributions of *R.*

As in Trials 1 and 2, we observed a rapid decrease in the average root detection rate during the first drought period for single and double drought treatment plants as compared to well-watered (Fig. 4(a), T-test, *p*<2.2e-16), whereby the average rate for single and double drought plants during the first drought period was 0.31 root detections per day, 64% less than that of well-watered plants during the same time period. Following the first drought, similar to water-limited plants in Trial 2, double drought plants in Trial 3 also temporarily exhibited increased detection rates relative to well-watered plants. The average daily root detection rate of double drought plants for a period leading into the second drought (8/18/19 to 8/28/19) was 0.87 roots per day - 40% greater than well-watered plants (T-test, *p*=2.98e-5). However, the detection rate of the double drought plants began to decrease once the second drought began. In contrast, single drought plants exhibited an increase in detection rates beginning at the same time as was observed for double drought plants and sustained increased growth to the end of the trial. The average daily root detection rate of single drought plants from 8/18/19 to 9/8/19 was 0.80 roots per day - 46% greater than well-watered plants (T-test, *p*=3.29e-9). In the time following the end of the first drought, we found that the cumulative detections of single drought plants approached those of well-watered plants (Fig. 4(a) inset). In contrast, and unlike in Trial 2, double drought plants in Trial 3 were unable to approach the average cumulative root detections of well-watered plants.

While not as pronounced as in Trial 2, we similarly observed that single and double drought plants exhibited greater rates of root detections at the deeper electrodes relative to well-watered plants following the first drought (Fig. 4(c)). The average daily root detection rate of single and double drought plants in the deeper half of the electrodes from 8/18/19 to 8/28/19 was 0.24 roots / day - 69% greater than well-watered plants (T-test, *p*=1.19e-8). Strikingly, even when the overall average detection rate of double drought plants was reduced to similar levels as well-watered plants following the initial increase in rate, double drought plants still exhibited more detections at the deeper electrodes than well-watered plants (Fig. 4(d)). The average daily root detection rate of double drought plants in the deeper half of electrodes from 8/28/19 to 9/8/19 was 0.12 roots / day - 79% greater than well-watered plants (T-test, *p*=0.0006). Consequently, the double drought root detection rates during this time period were also marked by fewer detections in the shallow region.

### The RootTracker identifies a wide array of root phenotypes present in maize

Previous analyses using shovelomics and gel-based growth media indicated that root phenotypes differ across a maize nested association mapping population (Zurek et al. 2015, Hauck et al. 2015). However, there is very little available data on root systems of commercial maize hybrids. To gain insight into the level of natural variation in root phenotypes among commercial maize hybrids, we used 1482 RTs to monitor root growth of 32 hybrids with high yield potential as well as six inbred lines. The trial was performed at Real Farm Research in Aurora, Nebraska in a loamy silt soil where corn and soybeans are typically grown. Inspection of root growth records over 55 days demonstrated that there were significant differences in root detection rates (Fig. 5, 1-way ANOVA, *F*(37) = 1.93, *p* = 0.0013). Two contrasting genotypes (H005 and H013) highlight these differences by both time and depth (Fig. 5b-d) These results provide evidence that a broad range of root phenotypes are present in the germplasm of modern maize hybrids.

**Figure 5.**
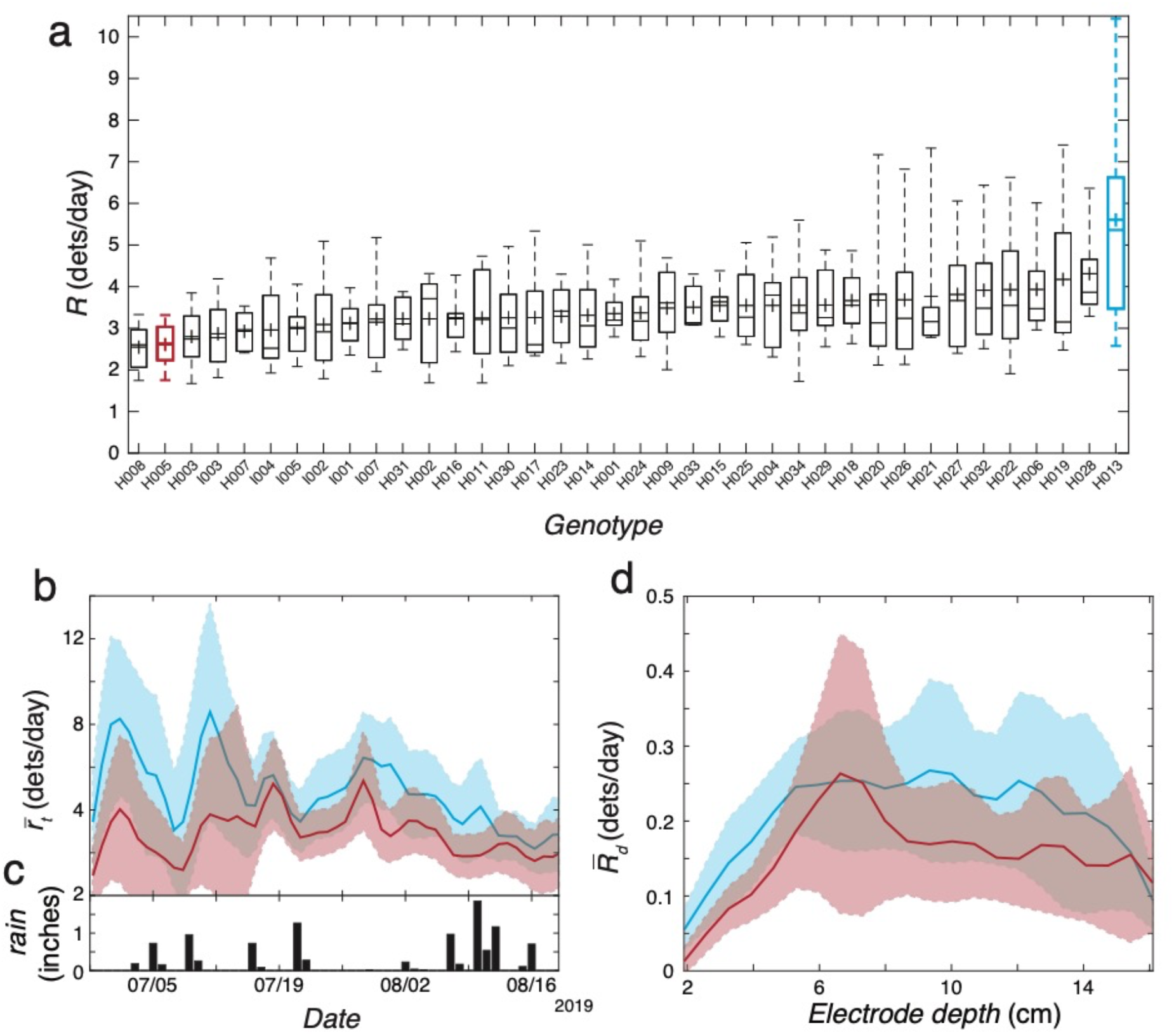
Trial 4: Comparison of maize genotypes in Midwest fields. (a) Box and whisker distributions of daily root detection rates time-averaged across the entire trial, *R,* separated by genotype. Top and bottom of box indicate 25 and 75 percentile of RTs, horizontal line in box is median, cross is mean, and whiskers are 9 and 91 percentiles. (b) Comparison of root detections between plants of genotype H005 (red) and H013 (blue) of (b) smoothed mean daily root detection rate over time, 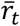, and (d) smoothed mean of the per-electrode-depth daily root detection rate, time-averaged across the entire trial, 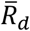. Shaded regions indicate ± 2 standard errors from the mean. (c) Inches of rain for each day received at Aurora, NE by station GHCND:US10hami004 (data provided by the NOAA/NCDC National Centers for Environmental Information, Asheville North Carolina from their website at ncdc.noaa.gov, Menne et al. 2020)

### RootTracker measurements correlate with shovelomics

To validate measurements made with the RT, we performed a ground-truth procedure by comparing RT detections with root mass measured by shovelomics (Trachsel et al. 2011). After Trial 1, roots were excavated, washed, imaged and analyzed according to the shovelomics protocol (see Methods).

To assess the accuracy of the RT, we used the excavated root images to measure the total pixels that were located in the same region of soil as the RT detectors would have been. Known dimensions of the RT, consistent placement of the root system in the frame of the image, and a known scale for pixels were used to identify this region (red rectangles in Fig. 6(b), see Methods for more details). We found a strong correlation between daily root detection rate time-averaged across the entire trial vs root pixels in the images (Fig. 6(c)), verifying that the RT platform is able to detect a similar root mass to that found by shovelomics. Compared to shovelomics, the RT has many advantages in that it is able to monitor root growth non-invasively, thus providing information as to root system growth over time and its response to environment factors, not just a measure of its final architecture.

**Figure 6.**
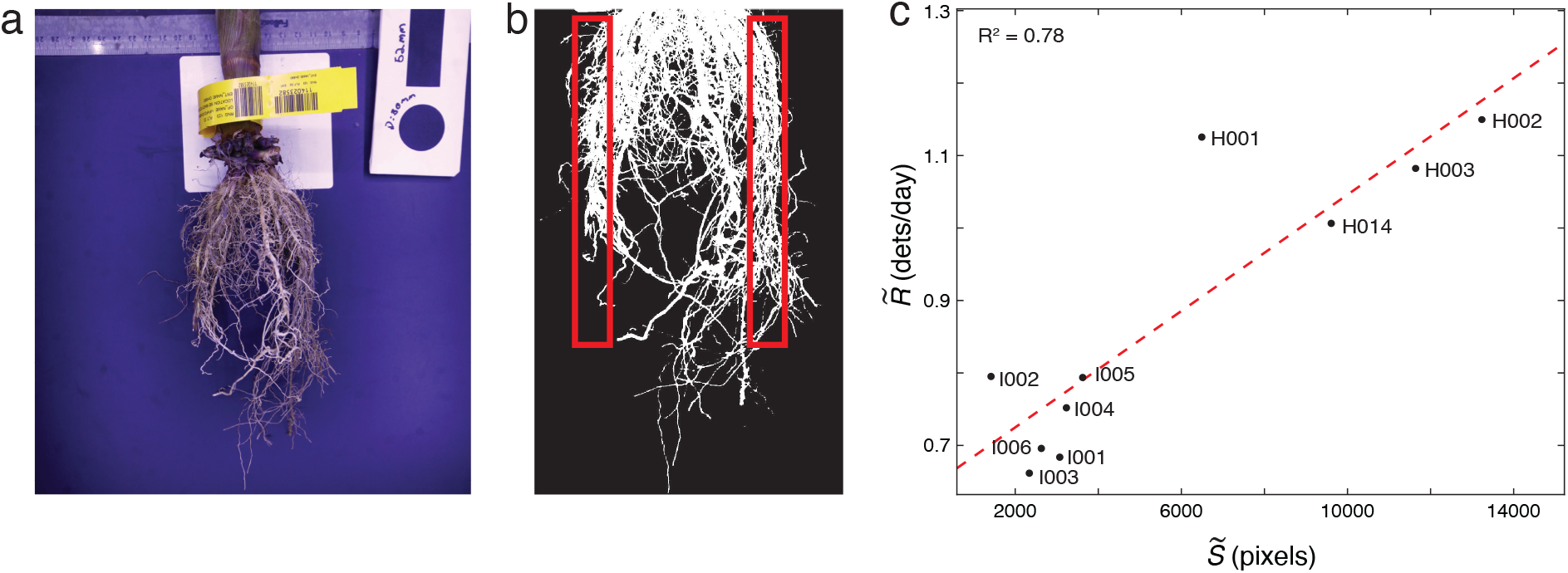
Shovelomics comparison in Trial 1. (a) Sample photograph of root system excavated from an RT. (b) Image segmenting for root pixels. Root pixels located in the region that would interact with RT detectors (red boxes) were counted for each plant. (c) Median daily root detection rate time-averaged across the entire trial, 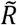, grouped by genotype, versus median shovelomics image root pixels, 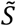, grouped by genotype. Correlation between RT detections and shovelomics root characterization: *R^2^* = 0.78.

## Discussion

Breeding based on direct measurement of a phenotype is superior to utilizing a secondary indicator, such as yield. To that end, the RT enables direct measurement of root system architecture and root system dynamics, phenotypes strongly connected to abiotic stress tolerance, especially tolerance to drought. Current methods to measure root growth in the field are either destructive (e.g. shovelomics) or sample only a small subset of roots (e.g. minirhizotrons). We have described a sensor-based technology able to monitor root growth over time in field conditions. In addition to providing a cost-effective and scalable platform for root phenotyping, communication of raw sensor data from remote locations is facilitated by relatively small file sizes (~1 kB per RT every 5 minutes) as compared to image-based modalities, and subsequent conversion of raw data to root detections occurs offline.

A strong correlation exists between plants with deeper roots and increased tolerance for water deficits (Li et al. 2019, Lynch 2013). However, there is little knowledge as to how roots *temporally* and spatially respond to water deficits in the field. Our results clearly show a rapid reduction in root growth when irrigation is shut off with a more dramatic response in the shallow soil strata. This suggests that maize plants can modulate root growth in response to small differences in soil moisture, opening the possibility of breeding plants with root systems optimized to respond to drought by growing deeper when there is a detected water deficit. The RT provides an opportunity to quantitatively characterize the response phenotypes in a wide array of different watering/drought scenarios (with variations in timing relative to plant development, duration, as well as severity) in both a controlled irrigation context as well as in the context of rain events. For example, daily root detection rates suggest that plants in Trial 4 exhibited growth rate fluctuations that coincided with rain events early in the trial (Fig. 5b-c).

The first few weeks of growth are important for the viability and robustness of row crops. An early exposure to abiotic stress known as “priming” has been shown to provide a measure of protection against later stresses in wheat. Our results suggest that the same may be true of maize. Root growth monitoring by the RT indicated that imposing an early water deficit resulted in maize plants with more and deeper roots later in the growing season. In Trials 2 and 3, we identified increased root growth subsequent to an early drought at a time coinciding with a second imposed drought. This increased growth also occurred in plants that were only subjected to a first priming drought. Furthermore, the single drought plants sustained increased growth for a longer period of time as compared with plants that were subjected to two droughts. This suggests that the increased growth rate was exclusively a priming effect resulting from the earlier water deficit, and the timing of the second drought in Trials 2 and 3 was coincidental relative to the timing of the observed augmented growth rates.

Monitoring cumulative detections over time indicated that the priming response may be a mechanism for plants to “catch-up” with root growth that would have occurred under well-watered conditions, and that root systems may target a specific root biomass. This would suggest that there is a genetic component to root biomass, which could be a target for breeding.

The differences in timing and duration of the drought treatments between Trials 2 and 3 and the resulting root growth highlight the responsiveness and adaptability of maize plants. Cumulative detections of the water-limited plants of Trial 2 approached those of well-watered plants in spite of the second drought. By contrast the double drought treatment plants in Trial 3 were unable to reach this target. The difference could be due to a longer first drought period (14 days) in Trial 3 as compared to Trial 2 (6 days), which constitutes a more severe early drought. The same growth reduction may have occurred in Trial 2 if the experiment had continued for a longer time period, since reduced root growth in the drought treatment plants was observed towards the end the trial (Fig. 3(c)).

Beyond aggregate root growth, monitoring roots with the RT revealed how plants can modulate the depth of new roots during and after imposed droughts. For example, towards the end of the second drought in Trial 3, when daily root growth of the double drought plants was similar to the well-watered plants, the double drought plants exhibited relatively greater deep roots and fewer shallow roots. This suggests a trade-off as to how to allocate new roots in search of water. Furthermore, while the single drought plants in Trial 3 primarily grew roots in the shallow soil strata towards the end of the experiment, they still exhibited greater root growth at the deeper electrodes than the well-watered plants. This suggests that plants may be able simultaneously to respond both to current environmental conditions as well as stresses that occurred at earlier growth stages.

It has been claimed that root systems for commercial hybrids must have been optimized during the intense selection for increased yield. Our data from a Midwestern field was not consistent with this claim, revealing differences in root growth patterns among commercial hybrids. This suggests that there remains a substantial amount of natural variation for root phenotypes in elite maize lines. A plausible reason is that most maize breeding has been performed under nutrient and water replete conditions presenting minimal selection pressure on root growth. The presence of alleles in the genome of elite cultivars for different root phenotypes provides an exciting opportunity to identify and breed for plants that are optimized for specific environments and that mitigate greenhouse gas emissions.

We describe a proof-of-concept use for the RT in identifying root phenotypes in maize. The platform can be adapted to other row crops, either in its present form or with modifications in its form factor. Moreover, it can be used both in the field and in controlled environment settings. Thus, the RT platform provides the opportunity to discover how roots grow in different soils and respond to different stimuli.

## Supporting information

Trial 1 RTs

Trial 2 RTs

Trial 3 RTs

Trial 4 RTs

## Acknowledgements

This work was supported by Advanced Research Projects Agency–Energy, grants DE-AR0000725 and DE-AR0000951. The trials were performed in conjunction with our collaborators: Massai Agricultural Services for Trials 1 and 2, Kearney Agricultural Research and Extension (KARE) Center for Trial 3, and Real Farm Research for Trial 4. Special thanks to Luis Liberon, Jose Manual De La Sotta, Chad Hill for their tireless efforts in helping with the trials. Special thanks to Daniel I. Goldman for early conversations in the use of capacitance touch sensing for detecting roots as well as the use of his x-ray equipment for preliminary 2-D real-time validation experiments.

