## Supplementary material for "Capturing in-field root system dynamics with the RootTracker": Methods

**RT Field Installation**

All fields were tilled prior to experiment installation to produce a soft seed bed and for ease of RT installation. Raised beds were formed for trials at Massai Agricultural Services in Rancagua, Chile and at the Kearney Agricultural Research and Extension (KARE) Center located in Parlier, California. Bed preparation was critical for improving water movement throughout the bed which increased the sensitivity of the RT units. Similarly, furrows between rows were created prior to trial installation at Real Farm Research (RFR) in Aurora, Nebraska. Standard fertilizer, weed and pest control for corn were used based on the recommendation of each cooperator.

Within each row, RTs were installed every two feet (every other plant, with the exception of Trial 2, where all plants spaced one foot apart within a row had an RT). After RTs were installed, seeds were sown by hand at two inches deep. In some trials, two seeds were sown and thinned to a single plant after emergence. In all trials, plants were spaced 1 ft apart within rows and between row spacing was 30 inches. See Supplemental Figures 1-4 for field layout maps and Supplemental Table 1 for a summary of RootTracker trials.

RootTracker installation is easiest in softer, tilled soil. Tool-free hand installation (i.e. pushing them into the ground by hand) is seldom possible. After soil preparation, RTs were set upright in the field. To minimize paddle bending during installation, we custom computer numerical control (CNC) fabricated 2 mm thick plastic guide rings to sleeve the paddles and keep them vertical during installation. The RTs were installed using a gas-powered fence post driver in conjunction with a custom-welded fence post hammer that mates with the RT’s center hole and the paddle tops. The process involves two operators, one operating the fence post driver as it rests on the hammer and another holding the hammer upright for proper vertical installation. As an RT is pushed into the ground, the guide ring presses up against the underside of the RT surface.

**Data logging and communication.**

Each RT records raw voltage measurements every 5 minutes. A radio module on the RT (the RFM69 HCW) communicates data upon measurement through one of several radio frequencies ranging from 902 to 924.5 MHz to solar powered base stations centrally located on the field. Each base station received signals from specific frequencies. Aside from distinct frequencies, the radio modules further filter radio traffic by signals transmitted on designated networks and nodes within that network. In each experiment, the RTs were pre-programmed to communicate on a unique node with an array of networks/frequencies. To minimize radio traffic interference among RTs, the base stations transmitted correction time delays to the RTs that ensured transmissions of all RTs within a network were evenly spaced within each 5-minute time window. The base stations compiled and compressed the received data in 15-minute increments and communicated via a cell modem to remote servers on Amazon Web Services (AWS). Data from AWS was regularly downloaded, parsed, processed, and analyzed on HFG servers located in Durham, NC.

**RT Tagging and Data Culling**

Each RT was a priori tagged with identifiers derived from its unique node/network/frequency assignment, which were used to record the field location of each RootTracker. RTs with no plant due to poor germination or premature death from physical damage (e.g. extreme weather or pest damage) were identified and excluded from all statistical calculations. The trials conducted in Rancagua, Chile also included a parallel technology development test, whereby two versions of the RootTracker were compared for detection accuracy. The newer design (Version 2, or V2), which comprised of 881 of the 1223 RTs in Chile, incorporated modified capacitance charging circuitry, and demonstrated superior shovelomics correlation as compared to the older design (Version 1, or V1) (Supplemental Figure 5). Consequently, only the V2 RT data from these experiments were utilized in our calculations (Figs 2, 3 and 6). Additionally, all RTs in subsequent trials (both Trial 3 and Trial 4) are V2 RTs.

**Drought trial irrigation methods**

The trials in Rancagua, Chile were irrigated using pressure-compensating drip tape laid on top of each raised bed and secured under the RTs. Drip tape in rows associated with the same treatment were connected to a main water line with valves placed to allow for treatment specific irrigation regimes by manually opening or closing valves. Irrigation time and duration were based on soil water holding capacity, estimated plant water demand during different plant growth stages, and average daily temperature, following standard corn irrigation methods of the cooperator. No rainfall events occurred throughout the course of these trials.

The trial at KARE was irrigated using drip irrigation. Drip lines were placed in the furrow bottoms between raised beds to avoid pooling on top of beds. The reasoning for this placement was that ponding of water in this sandy loam soil can result in a silty hard crust on the soil surface. Total weekly crop water demand was estimated using the previous week’s crop evapotranspiration (ETc). ETc was estimated using weather data from the California CIMIS station located in Parlier and estimated plant water demand for different maize plant growth stages. Applying a system pressure of 10 psi, the average drip emitter water output was checked at intervals to assess relative uniformity of drip water application amounts. This was estimated by measuring the amount of water emitted over a set time period for 12 to 16 emitters (varied at different times), with the emitters evaluated distributed across the entire field. Given this average volumetric output, the average water application rate of the system was calculated to be 0.19 inches/hour. This rate was used in conjunction with the estimated ETc to determine the total irrigation time per week. For example, according to the CIMIS station data and crop demand, ETc for the week of 8/12/19-8/18/19 was estimated to be 1.99 inches. Thus, given a watering rate of 0.19 inches/hour, the recommended watering time for the following week was 10.5 hours. The total irrigation time for a given week was distributed across multiple days to allow the soil to dry sufficiently between irrigation events and aid our ability to walk in the field to collect above ground plant measurements and observations.

**Chile shovelomics methods**

In January of 2019, all RTs in drought trial 1 were excavated by hand using shovels, keeping the roots intact within a one foot diameter and 8 inch depth around the RT. Plants were cut above the brace roots and roots were carefully removed from RTs, keeping track of the associated RT, field location, genotype, and treatment. Roots were gently washed in large bins of water with mild detergent. After washing, roots were laid out to air dry. Dry roots were photographed in a photo area constructed to capture images with consistent lighting, focus, and distance from the roots. The imaging setup also provided consistent contrast between the root system and backdrop, allowing for accurate identification of image pixels containing root matter (Fig. 5(a)).

The goal of the shovelomics image analysis was to approximate the amount of root matter in close proximity to the region of the RT where paddle electrodes were located. For each image, a threshold pixel intensity was used to identify and count all root pixels, $S$, located in the region of paddle electrodes. Known dimensions of the RT, consistent placement of the root system in the image frame, and a known pixel scale were used to identify this region (red rectangle in Fig. 5(b)).

Shovelomics root pixel calculations or each RT were compared with $R$, the daily root detection rate, time-averaged across the entire trial period calculated for each RT. Correlations were generated via grouping $R$ and $S$ by genotype and calculating respective median values $\tilde{R}$ and $\tilde{S}$.

**Root detection calculations**

RootTracker detections are determined from processed raw voltage signals. Using direct current charging, each electrode is charged while all other sensors are grounded, and voltage at that electrode is measured multiple times with different charge times. Assuming a simple parallel resistor-capacitor circuit between a charged electrode and surrounding grounded electrodes, an estimate of capacitance and resistance is calculated using measured voltages.

$Res= \frac{{Res}_{VD}V_{2}}{V_{s}-V_{2}}$	Eq. 1

$Res$ is resistance; ${Res}_{VD}$ is the voltage divider resistance; $V_{2}$ is the measured voltage of the electrode for a time sufficiently long to assume no capacitance effect on voltage measurement, and $V_{s}$ is the source voltage.

$Cap= \frac{-T_{C}(Res+Res_{VD})}{Res\cdot Res_{VD}ln(1-{{(V}_{1}}/{V_{s}})(Res+ Res_{VD})/Res)}$	Eq.2

$Cap$ is capacitance, $T_{C}$ is the short time used to charge the electrode, $V_{1}$ is the measured voltage of the electrode when charge for $T_{C}$ time. We have identified signature fluctuations in the resistance/capacitance space that indicate root growth activity near the sensors once signal changes have been normalized across all electrodes; thus, fluctuations are evaluated to detect roots.

The detection algorithm masks portions of the data deemed unreliable, such as short periods of dramatic rapid signal changes that indicate the moment of a watering event like rain or irrigation, or voltage measurements so low (due to locally saturated water) as to cause low resolution data and consequently unreliable R-C calculations. Additionally, at times, issues with the base station or individual root trackers result in temporary down time whereby RT data is simply unavailable for analysis. Collectively, this missing data affects a RootTracker’s daily fractional uptime, $U_{t}$. Thus, to calculate a RootTracker’s daily root detection rate, $r_{t}$, we normalize by the RT’s $U_{t}$ for that day:

$r_{t}= \frac{n_{t}}{U_{t}}$	Eq.3

where $n_{t}$ is the number roots detected for an RT in a given day, $t$. Days with $U_{t}\leq0.04$ were treated as $U_{t}=0$ and were ignored to avoid unrealistic values of $r_{t}$ due to limited available data. We calculated daily growth rate by day at a specific electrode depth, $r_{td}$, by aggregating all detections of an RT recorded at a specific electrode depth, $d$, (from any of the RT’s 12 paddles), $n_{td}$, and normalizing by the relative amount of data available at that depth for that day $U_{td}$.

$r_{td}= \frac{n_{td}}{U_{td}}$	Eq.4

We applied a low-pass Butterworth filter to plots of mean growth rates over time, $\bar{r}_{t}$ (and their respective 2x standard errors) with a normalized cutoff frequency of 0.4. We similarly applied a low pass filter to heatmaps of mean growth by depth and time, $\bar{r}_{td}$, filtering first by depth, then by time, both passes with cutoff filters of 0.4. The same filtering was applied to plots of mean time-averaged growth rates by depth, $\bar{R}_{d}$ (and their respective 2x standard errors).To calculate the cumulative roots detected over time, $c_{t}$we integrate aggregate $r_{t}$ over time:

$c_{t}= \sum_{i=0}^{i=t} r_{i}$	Eq.5

We calculate time-averaged growth rates as follows:

$R= \frac{\sum_{t=a}^{t=b} r_{t}}{b-a}$	Eq.6

where $a$ and $b$ are the start and end days of the time-averaged period, respectively. Similarly, we calculate time-averaged growth rates at specific electrode depths as follows:

$R= \frac{\sum_{t=a}^{t=b} r_{td}}{b-a}$	Eq.7

In box and whisker plots of $R$ for different RTs of a specific group, such as seen Fig. 2(d) or 5(a), we excluded RTs from the distribution that had minimal data available for their time average. Specifically, we removed RTs where more than 50% of the days of the time-averaging period had no data available ($U_{t}=0$) or where the span of days with data available for calculation was less than 80% of the period date range.
