## Supplemental Figures for "Capturing in-field root system dynamics with the RootTracker"

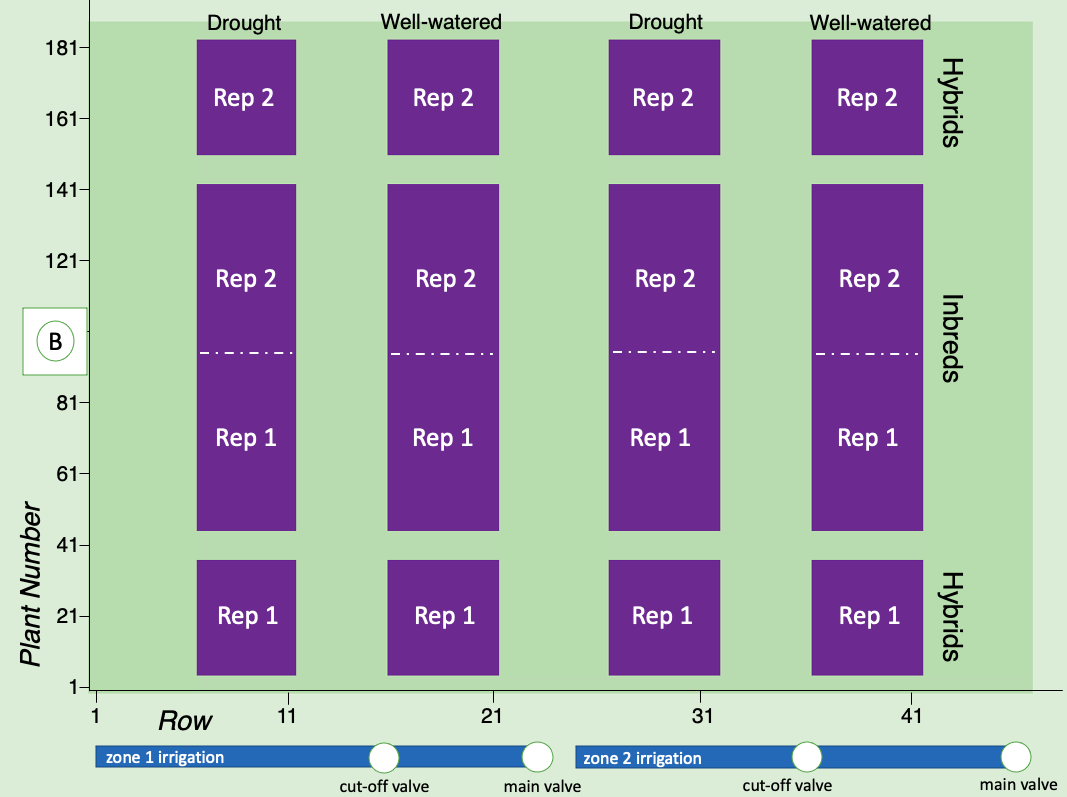

Supplemental Figure 1. Trial 1 field map (Massai Agricultural Services, Rancagua, Chile, 2018-19). Rows were spaced 30” apart and plants within rows were 1’ apart. The base stations (B) were located to the left of the field half-way down the rows. Two irrigation treatments (drought and well-watered) were applied in blocks 5 rows wide and 173 feet long (purple), separated by 5 or 6 rows of border plants (dark green). The experiment included two reps of each treatment, and 10 genotypes (4 hybrids and 6 inbred lines) were tested across each treatment block in two reps. To minimize neighboring canopy and treatment effects, the field was laid out in a split plot design in with hybrids and inbreds blocked within drought and well-watered treatments blocks. Genotypes were randomized within blocking factors and planted in plots 8 plants long by 5 rows wide. Drip lines were attached to one of two main water lines (irrigation zones 1 and 2). Cutoff valves allowed for a drought and a well-watered section along each main water line. Version 2 RTs were primarily located in the right two treatment blocks, and version 1 RTs were installed primarily in the left two treatment blocks. See supplementary file “RTs_T1.csv” for a list of RootTrackers and their corresponding treatment, genotype, hardware version and location on the field.

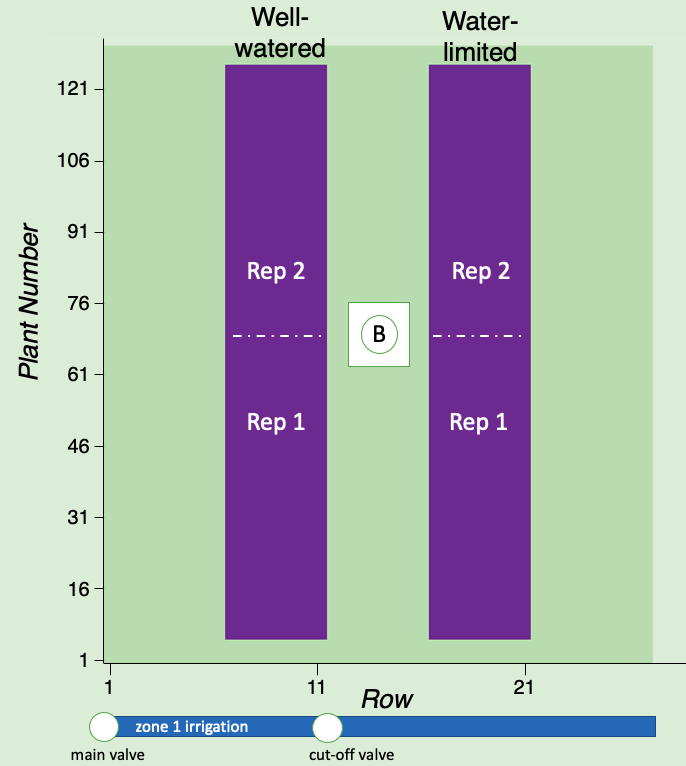

Supplemental Figure 2. Trial 2 field map (Massai Agricultural Services, Rancagua, Chile, 2019). Rows were spaced 30” apart and plants within rows were 1’ apart. The base stations were located at the center of the field. Treatments (well-watered and water-limited) were applied in separate blocks 5 rows wide and 120 feet long (purple), separated by 6 rows of border plants (dark green). All plants in treatment rows (purple) were in RTs. Drip lines were attached to a main water line (irrigation zone 1) with a cutoff valve that allowed for separate well-watered and water-limited sections. The experiment included 12 hybrid genotypes. Genotypes were randomized within treatment blocks, planted in plots 5 plants long by 5 rows wide, and tested across each treatment block in two reps. Version 2 RTs were primarily installed at bottom half of the water-limited treatment block. See supplementary file “RTs_T2.csv” for a list of RootTrackers and their corresponding, treatment, genotype, hardware version and location on the field.

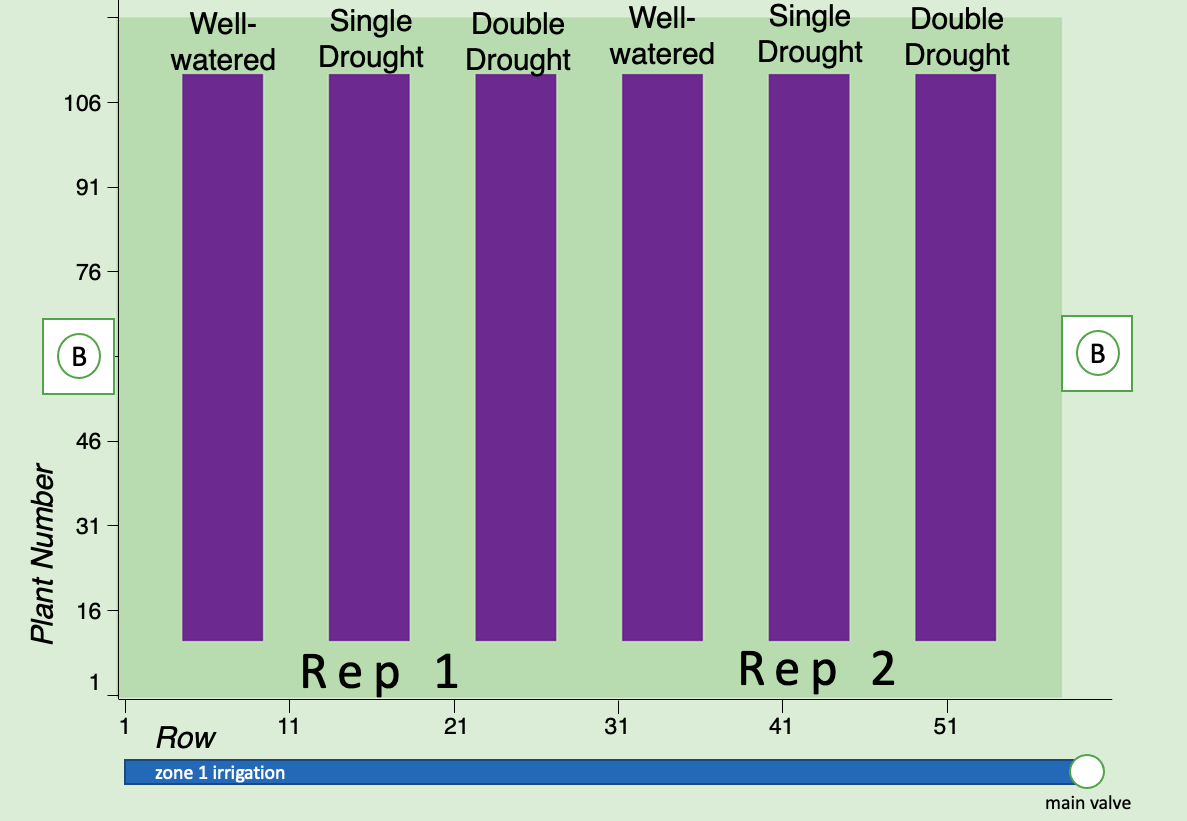

Supplemental Figure 3. Trial 3 field map (Kearney Agricultural Research and Extension (KARE) Center located in Parlier, California, 2019). Rows were spaced 30” apart and plants within rows were 1’ apart. The base stations were located half-way down the rows on either side of the field. Treatments (well-watered, single drought, and double drought) were applied in separate blocks 5 rows wide and 99 feet long (purple), separated by 4 rows of border plants (dark green). Treatment rows (purple) contained RTs for every other plant. Drip lines were attached to a main water line (irrigation zone 1) with a cutoff valve that allowed for separate well-watered and water-limited sections. Each drip line was individually controlled by a manual valve. The experiment included 10 hybrid genotypes. Genotypes were randomized within treatment blocks, planted in plots 10 plants long by 5 rows wide, and tested across each treatment block in two reps. See supplementary file “RTs_T3.csv” for a list of RootTrackers and their corresponding, treatment, genotype and location on the field.

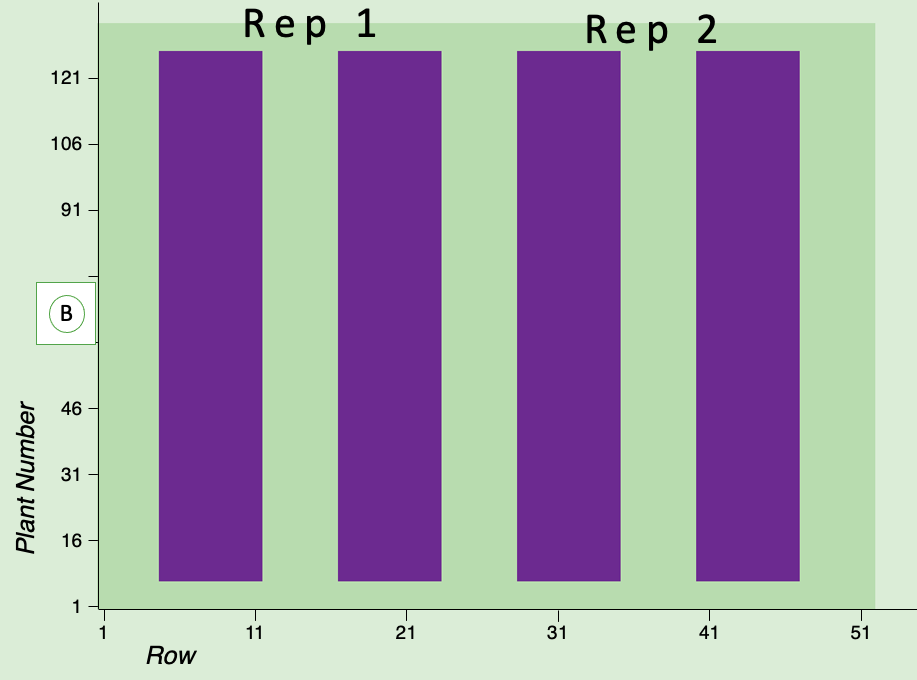

Supplemental Figure 4. Trial 4 field map (Real Farm Research, Aurora, Nebraska, 2019). Rows were spaced 30” apart and plants within rows were 1’ apart. The base stations were located to the left of the field half-way down the rows. Experimental blocks of the field (purple) were 7 rows wide by 114 feet long and were surrounded by 4 or 5 rows of border plants (dark green). Rows in the experimental blocks contained RTs every other plant (i.e. spaced 2 ft apart). Experimental blocks were 7 rows wide and 113 plants long. The experiment included 25 hybrid and 13 inbred genotypes tested in two reps, where the left two experimental blocks contained rep 1 and the right two contained rep 2. To minimize neighboring canopy and treatment effects, hybrids and inbreds were blocked separately. Genotypes were randomized within inbreds and hybrids in both reps and were planted in plots 6 plants long by 7 rows wide. See supplementary file “RTs_T4.csv” for a list of RootTrackers and their corresponding genotype and location on the field.

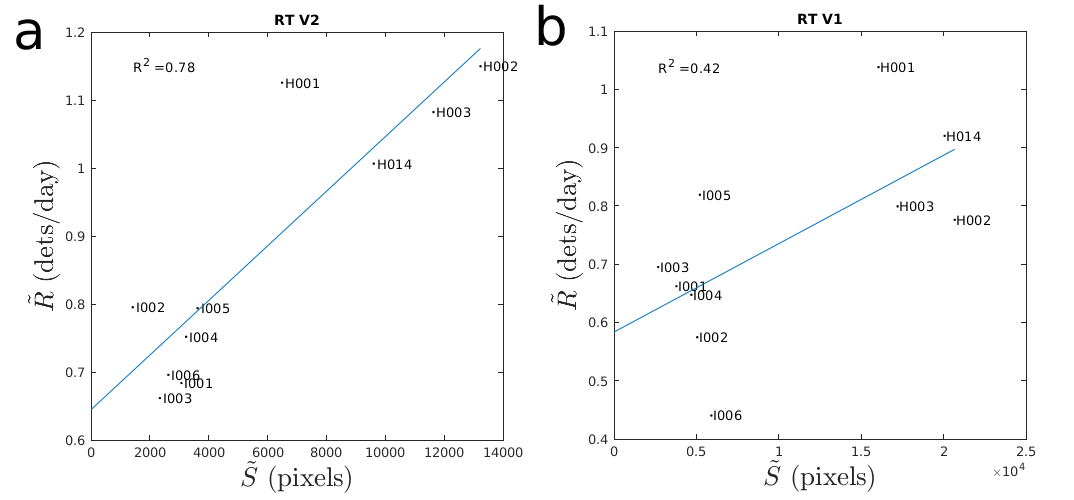

Supplemental Figure 5. Shovelomics comparison of different RT versions. Median daily root detection rate time-averaged across the entire trial, $\tilde{R}$, grouped by genotype, versus median shovelomics image root pixels, $\tilde{S}$, grouped by genotype for (a) V2 RTs vs (b) V1 RTs. The primary difference between the two hardware versions is a change in V2 to the resistance in the voltage divider of the charging circuit allowing for great signal sensitivity in wet soils.

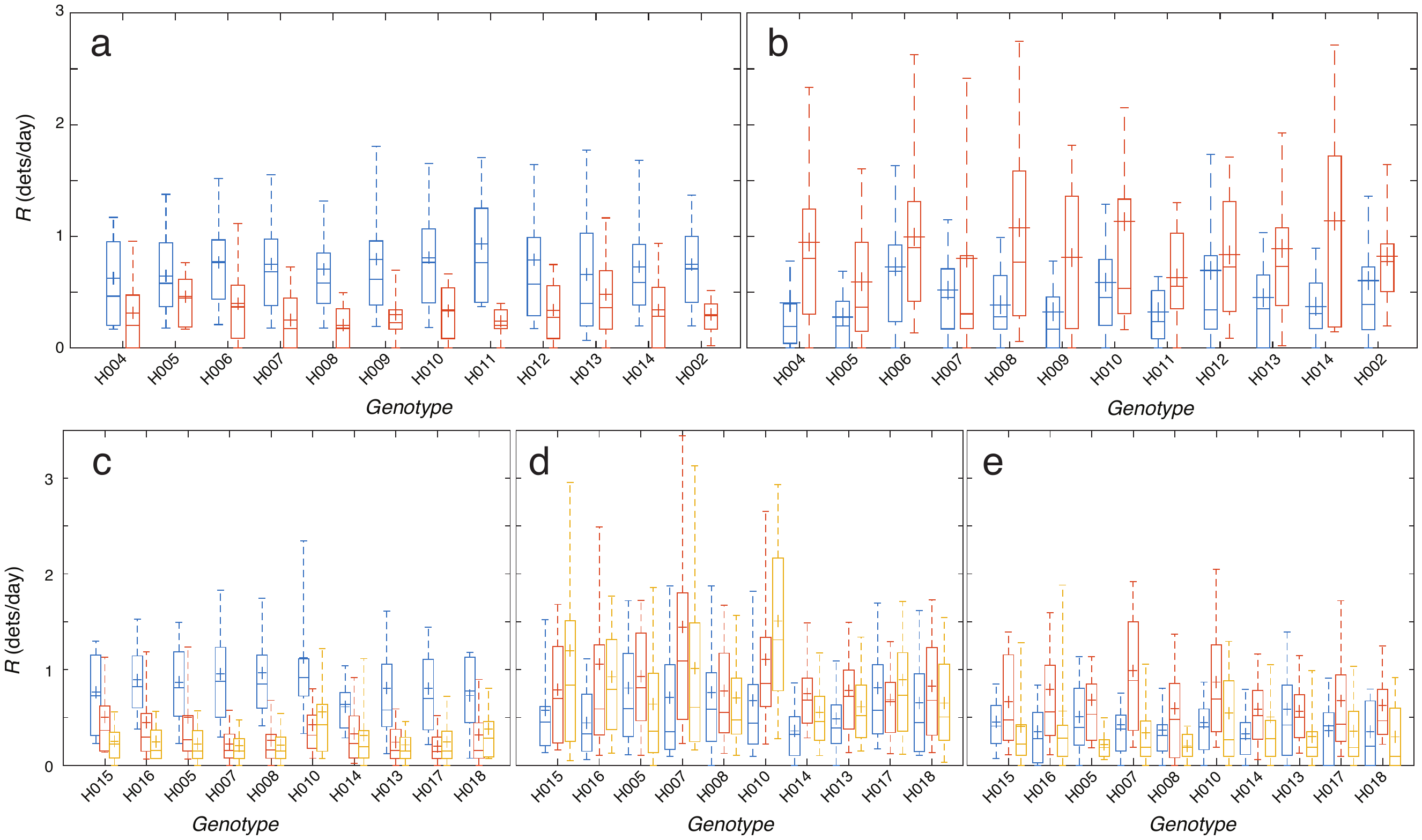

Supplemental Figure 6. Box and whisker distributions of time-averaged daily root detection rates, $R$, separated by genotype and treatment, time-averaged during (a) a period (2/16/2019 - 2/22/2019) overlapping with the early drought in Trial 2, (b) part of the late drought in Trial 2 (3/15/19 - 3/22/19), (c) the early drought in Trial 3 (7/26/2019 - 8/9/2019), (d) a period (8/18/2019 - 8/28/2019) overlapping with the beginning of the second drought in Trial 3, and (e) a period (8/28/2019 - 9/8/2019) overlapping with the end of the second drought in Trial 3. For Trial 2 (a,b), the color code is as follows: well-watered: blue; water-limited: orange. For Trial 3 (c-e), the color code is as follows: well-watered: blue; single drought: orange; double drought: yellow.

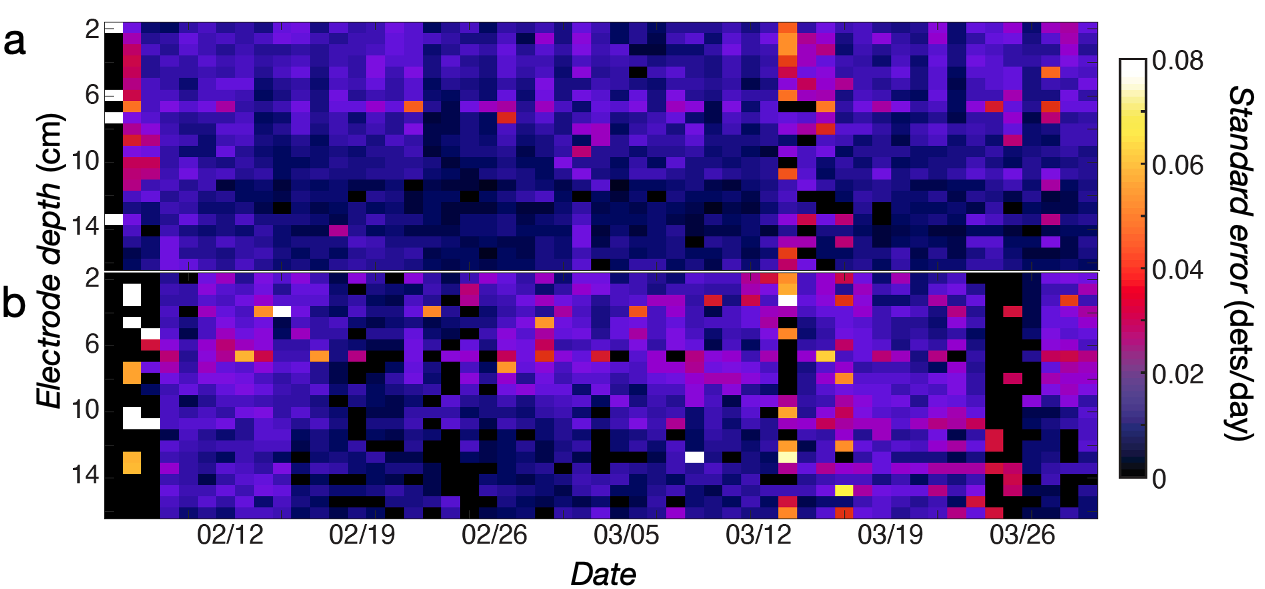

Supplemental Figure 7. Heatmaps of standard error for mean daily root detection rate over time and electrode depth, $\bar{r}_{td}$ for (a) well-watered plants, and (b) water-limited plants in Trial 2.

| Trial Name | Location | Trial Period (planting date to end) | Key Time Periods | Treatments | Total # of RTs |
| --- | --- | --- | --- | --- | --- |
| Trial 1 | Rancagua, Chile | 11/29/2018-1/20/2019 | drought \| 1/4/2019-1/20/2019 | well-watered; drought | 1223 (342 V1 RTs, 881 V2 RTs) |
| Trial 2 | Rancagua, Chile | 2/5/2019-3/30/2019 | early drought \| 2/14/2019-2/20/2019; late drought \| 3/12/2019-3/23/2019 | well-watered; water-limited | 1154 (433 V1 RTs, 721 V2 RTs) |
| Trial 3 | Parlier, California | 7/10/2019-9/8/2019 | early drought \| 7/26/2019-8/9/2019; late drought \| 8/23/2019-9/8/2019 | well-watered; single drought; double drought | 1457 |
| Trial 4 | Aurora, Nebraska | 6/28/2019-8/22/2019 | N/A | N/A | 1482 |

Supplemental Table 1. Summary of RootTracker trials.
